# Neurophysiological indices of audiovisual speech integration are enhanced at the phonetic level for speech in noise

**DOI:** 10.1101/2020.04.18.048124

**Authors:** Aisling E. O’Sullivan, Michael J. Crosse, Giovanni M. Di Liberto, Alain de Cheveigné, Edmund C. Lalor

## Abstract

Seeing a speaker’s face benefits speech comprehension, especially in challenging listening conditions. This perceptual benefit is thought to stem from the neural integration of visual and auditory speech at multiple stages of processing, whereby movement of a speaker’s face provides temporal cues to auditory cortex, and articulatory information from the speaker’s mouth can aid recognizing specific linguistic units (e.g., phonemes, syllables). However it remains unclear how the integration of these cues varies as a function of listening conditions. Here we sought to provide insight on these questions by examining EEG responses to natural audiovisual, audio, and visual speech in quiet and in noise. Specifically, we represented our speech stimuli in terms of their spectrograms and their phonetic features, and then quantified the strength of the encoding of those features in the EEG using canonical correlation analysis. The encoding of both spectrotemporal and phonetic features was shown to be more robust in audiovisual speech responses then what would have been expected from the summation of the audio and visual speech responses, consistent with the literature on multisensory integration. Furthermore, the strength of this multisensory enhancement was more pronounced at the level of phonetic processing for speech in noise relative to speech in quiet, indicating that listeners rely more on articulatory details from visual speech in challenging listening conditions. These findings support the notion that the integration of audio and visual speech is a flexible, multistage process that adapts to optimize comprehension based on the current listening conditions.

**Significance Statement:** During conversation, visual cues impact our perception of speech. Integration of auditory and visual speech is thought to occur at multiple stages of speech processing and vary flexibly depending on the listening conditions. Here we examine audiovisual integration at two stages of speech processing using the speech spectrogram and a phonetic representation, and test how audiovisual integration adapts to degraded listening conditions. We find significant integration at both of these stages regardless of listening conditions, and when the speech is noisy, we find enhanced integration at the phonetic stage of processing. These findings provide support for the multistage integration framework and demonstrate its flexibility in terms of a greater reliance on visual articulatory information in challenging listening conditions.

## Introduction

One prominent theory of speech perception is that speech is processed in a series of computational steps that follow a hierarchal structure, with different cortical regions being specialised for processing different speech features (Scott and Johnsrude, 2003; Hickok and Poeppel, 2007; DeWitt and Rauschecker, 2012). One key question is how visual input influences processing within this hierarchy.

Behavioral studies have shown that seeing the face of a speaker improves speech comprehension (Sumby and Pollack, 1954; Grant and Seitz, 2000; Ross et al., 2007). This behavioral advantage is thought to derive from two concurrent processing modes: a correlated mode, whereby visual speech dynamics provide information on auditory speech dynamics; and a complementary mode, where visual speech provides information on the articulatory patterns generating the auditory speech (Campbell, 2008). It seems plausible that the information provided by these two modes would influence levels of the auditory hierarchy differently. Indeed, this idea aligns well with a growing body of evidence indicating that audiovisual (AV) speech integration likely occurs over multiple stages (Schwartz et al., 2004; van Wassenhove et al., 2005; Eskelund et al., 2011; Peelle and Sommers, 2015; Baart et al., 2014). One recent perspective (Peelle and Sommers, 2015) suggests that these stages could include an early stage, where visual speech provides temporal cues about the acoustic signal (correlated mode), and a later stage, where visual cues that convey place and manner of articulation could be integrated with acoustic information to constrain lexical selection (complementary mode). Such early-stage integration could be mediated by direct projections from visual cortex that dynamically affect the sensitivity of auditory cortex (Grant and Seitz, 2000; Okada et al., 2013; Tye-Murray et al., 2011; Calvert et al., 1997), whereas for later-stage integration, articulatory visual cues could be combined with acoustic information in supramodal regions such as the STS (Beauchamp et al., 2004; Kayser and Logothetis, 2009; Zhu and Beauchamp, 2017; Karas et al., 2019).

While the evidence supporting multiple stages of audiovisual speech integration is compelling, there are several ways in which this multistage model needs to be further developed. First, much of the supporting evidence has been based on experiments involving simple (and often illusory) syllabic stimuli or short segments of speech. This has been very valuable, but it also seems insufficient to fully explore how a correlated mode of audiovisual integration might derive from dynamic visual cues impacting auditory cortical processing. Testing the model with natural speech will be necessary (Theunissen et al., 2000; Hamilton and Huth, 2018). Second, directly indexing neurophysiological representations of different acoustic and articulatory features will be important for validating and further refining the key idea that integration happens at different stages. And third, it will be important to test the hypothesis that this multistage model is flexible, whereby the relative strength of integration effects at different stages might depend on the listening conditions and the availability of visual information.

These are the goals of the present manuscript. In particular, we aim to build on recent work that examined how visual speech affected neural indices of audio speech dynamics using naturalistic stimuli (Luo et al., 2010; Golumbic et al., 2013; Crosse et al., 2015a; Crosse et al., 2016b). We aim to do so by incorporating ideas from recent research showing that EEG and MEG are sensitive not just to the acoustics of speech, but also to the processing of speech at the level of phonemes (Di Liberto et al., 2015; Brodbeck et al., 2018; Khalighinejad et al., 2017). This will allow us to derive indices of dynamic natural speech processing at different hierarchical levels and to test the idea that audiovisual speech integration occurs at these different levels, in line with the multistage model (Peelle and Sommers, 2015). Finally, we also aim to test the hypothesis that, in the presence of background noise, there will be a relative increase in the strength of AV integration effects in EEG measures of phoneme-level encoding, reflecting an increased reliance on articulatory information when speech is noisy. To do all this, we introduce a new framework for indexing the electrophysiology of audiovisual speech integration based on canonical correlation analysis (CCA).

## Methods

The EEG data analyzed here were collected as part of previous studies published by Crosse et al. (2015a; 2016b).

### Participants

Twenty-one native English speakers (eight females; age range: 19-37 years) participated in the speech in quiet experiment. Twenty-one different participants (six females; age range: 21-35) took part in the speech in noise experiment. Written informed consent was obtained from each participant beforehand. All participants were native English speakers, were free of neurological diseases, had self-reported normal hearing, and had normal or corrected-to-normal vision. The experiment was approved by the Ethics Committee of the Health Sciences Faculty at Trinity College Dublin, Ireland.

### Stimuli and procedure

The speech stimuli were drawn from a collection of videos featuring a trained male speaker. The videos consisted of the speaker’s head, shoulders, and chest, centered in the frame. The speech was conversational-like and continuous, with no prolonged pauses between sentences. Fifteen 60 s videos were rendered into 1280×720 pixel movies in VideoPad Video Editor (NCH Software). Each video had a frame rate of 30 frames per second, and the soundtracks were sampled at 48 kHz with 16-bit resolution. The intensity of each soundtrack, measured by root mean square, was normalized in MATLAB (MathWorks). For the speech in noise experiment, the soundtracks were additionally mixed with spectrally matched stationary noise to ensure consistent masking across stimuli (Ding and Simon, 2013; Ding et al., 2013) with SNR of −9 dB. The noise stimuli were generated in MATLAB using a 50th-order forward linear predictive model estimated from the original speech recording. Prediction order was calculated based on the sampling rate of the soundtracks (Parsons, 1987).

In both experiments, stimulus presentation and data recording took place in a dark sound attenuated room with participants seated at a distance of 70 cm from the visual display. Visual stimuli were presented on a 19 inch CRT monitor operating at a refresh rate of 60 Hz. Audio stimuli were presented diotically through Sennheiser HD650 headphones at a comfortable level of ~65 dB. Stimulus presentation was controlled using Presentation software (Neurobehavioral Systems). For the speech in quiet experiment each of the 15 speech passages was presented seven times, each time as part of a different experimental condition. Presentation order was randomized across conditions, within participants. While the original experiment had seven conditions, here we focus only on three conditions audio-only (A), visual-only (V) and congruent audio-visual (AVc). For the speech in noise experiment, however, there were only 3 conditions (A, V and AV) and so the passages were ordered 1-15 and presented 3 times with the condition from trial-to-trial randomized. This was to ensure that each speech passage could not be repeated in another modality within 15 trials of the preceding one. Participants were instructed to fixate on either the speaker’s mouth (V, AVc) or a gray crosshair (A) and to minimize eye blinking and all other motor activity during recording.

For both experiments participants were required to respond to target words via button press. Before each trial, a target word was displayed on the monitor until the participant was ready to begin. All target words were detectable in the auditory modality except during the V condition, where they were only visually detectable. A target word was deemed have been correctly detected if subjects responded by button press within 0–2 seconds after target word onset. In addition to detecting target words, participants in the speech-in-noise experiment were required to rate subjectively the intelligibility of the speech stimuli at the end of each 60-s trial. Intelligibility was rated as a percentage of the total words understood using a 10-point scale (0–10%, 10–20%, … 90–100%).

### EEG acquisition and preprocessing

The EEG data were recorded using an ActiveTwo system (BioSemi) from 128 scalp electrodes and two mastoid electrodes. The data were low-pass filtered on-line below 134 Hz and digitized at a rate of 512 Hz. Triggers indicating the start of each trial were recorded along with the EEG. Subsequent preprocessing was conducted off-line in MATLAB; the data were detrended by subtracting a 50th-order polynomial fit using a robust detrending routine (de Cheveigné and Arzounian, 2018). The data were then bandpass filtered using second-order, zero phase-shift Butterworth filters between 0.3-30 Hz, downsampled to 64 Hz, and rereferenced to the average of the mastoid channels. Channels contaminated by noise were recalculated by spline-interpolating the surrounding clean channels in EEGLAB (Delorme and Makeig, 2004).

### Indexing neurophysiological speech processing at different hierarchical levels

Because our aim was to examine how visual information affects the neural processing of auditory speech at different hierarchical levels, we need to derive separable EEG indices of processing at these levels. To do this, we followed work from Di Liberto et al. (2015) who modeled EEG responses to speech in terms of different representations of that speech. Specifically, they showed that EEG responses to speech were better predicted using a representation of speech that combined both its low-level acoustics (i.e., its spectrogram) and a categorical representation of its phonetic features. The underlying idea is that EEG responses might reflect the activity of neuronal populations in auditory cortex that are sensitive to spectrotemporal acoustic fluctuations and of neuronal populations in association cortices (e.g., the superior temporal gyrus) that may be invariant to spectrotemporal differences between utterances of the same phoneme and, instead, are sensitive to that phoneme category itself. As such, for the present study, we calculated two different representations of the acoustic speech signal.

1. **Spectrogram:** This was obtained by first filtering the speech stimulus into 16 frequency bands between 80 and 3000 Hz using a compressive gammachirp auditory filter bank that models the auditory periphery (Irino and Patterson, 2006). Then the amplitude envelope for each frequency band was calculated using the Hilbert transform, resulting in 16 narrow band envelopes forming the spectrogram representation.
2. **Phonetic features:** This representation was computed using the Prosodylab-Aligner (Gorman et al., 2011) which, given a speech file and the corresponding textual orthographical transcription, automatically partitions each word into phonemes from the American English International Phonetic Alphabet (IPA) and performs forced-alignment (Yuan and Liberman, 2008), returning the starting and ending time-points for each phoneme. Manual checking of the alignment was then carried out and any errors corrected. This information was then converted into a multivariate time-series that formed a binary array, where there is a one representing the onset and duration of each phoneme and zeros everywhere else. To describe the articulatory and acoustic properties of each phoneme a 19-dimensional phonetic feature representation was formed using the mapping defined by (Mesgarani et al., 2014; Chomsky and Halle, 1968). This involves mapping each phoneme (e.g., /b/) into a set of phonetic features (e.g., bilabial, plosive, voiced, obstruent) and results in a phonetic feature matrix of ones and zeros that is of dimension 19 (which is the number of phonetic features) by time.

### Canonical correlation analysis

We wished to see how these different speech representations might be reflected in EEG activity. Previous related research has relied on a regression-based approach that aims to reconstruct an estimate of some univariate feature of the speech stimulus (e.g., its amplitude envelope) from multivariate EEG responses (Crosse et al., 2015a; Crosse et al., 2016b). However, because we have multivariate speech representations, we sought to use a method based on canonical correlation analysis (CCA, de Cheveigne et al., 2018; Hotelling, 1936) which was implemented using the NoiseTools toolbox (http://audition.ens.fr/adc/NoiseTools/).

CCA works by rotating two given sets of multidimensional data into a common space in which they are maximally correlated. This linear transformation is based on finding a set of basis vectors for each of the given data sets such that the correlation between the variables, when they are projected on these basis vectors, is mutually maximized. In our case, our two data sets are the multidimensional stimulus representation, *X*(*t*), of size *T* × *J*_1_, where *T* is time and *J*_1_ is the number of features in that representation (*J*_1_ = 16 frequency bands of a spectrogram, or *J*_1_ = 19 phonetic features), and an EEG data matrix *Y*(*t*) of size *T* × *J*_2_, where *T* is time and *J*_2_ = *n × τ*, where *n* is the number of EEG channels (128) and *τ* is the number of time-lags. The reason for using multiple time lags is to allow for the fact that a change in the stimulus impacts the EEG at several subsequent time lags. In our analysis we included time-lags from 0-500 ms, which at a sampling rate of 64 Hz resulted in 32 time-lags. For these two data matrices, CCA produces transform matrices *A* and *B* of sizes *J*_1_ × *J*_0_ and *J*_2_ × *J*_0_ respectively, where *J*_0_ is at most equal to the smaller of *J*_1_ and *J*_2_. The optimization problem for CCA is formulated as a generalized eigenproblem with the objective function:

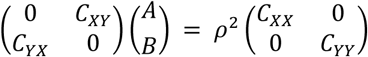

Where *C*_*XY*_ is the covariance of the two datasets X and Y and *C*_*XX*_ and *C*_*YY*_ are the autocovariances, and *ρ* is the components correlation. Ridge regularization can be performed on the neural data to prevent overfitting in CCA as follows (Vinod, 1976; Cruz-Cano and Lee, 2014; Leurgans et al., 1993; Bilenko and Gallant, 2016):

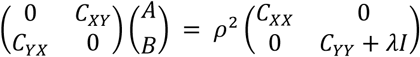

The rotation matrices (*A* and *B*) are learned on all trials except one and are then applied to the left-out data which produces canonical components (CC’s) for both the stimulus representation and the EEG using the following equation:

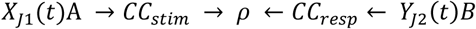

The rotation weights *A* and *B* are trained to find what stimulus features influence the EEG and what aspects of the EEG are responsive to the stimulus, respectively, in order to maximize the correlation between the two multivariate signals. When the rotation weights are applied to the left-out data we get the canonical components of the stimulus (*CC*_*stim*_) and of the response data (*CC*_*resp*_). The first pair of canonical components define the linear combinations of each data set with the highest possible correlation. The next pair of CCs are the most highly correlated combinations orthogonal to the first, and so-on (de Cheveigne et al., 2018).

### Indexing multisensory integration using CCA

We wished to use CCA to identify any neural indices of multisensory integration during the AV condition beyond what might be expected from the unisensory processing of audio and visual speech. We sought to do this by modelling the encoding of the speech representations in the audio-only (A) and visual-only (V) EEG data and then investigating if there is some difference in the speech-related activity in the AV EEG data which is not present in either of the unisensory conditions. In other words, and in line with a long history of multisensory research, we sought to compare AV EEG responses to A+V EEG responses using CCA and to attribute any difference (i.e., AV – (A+V)) to multisensory processing.

To implement this, we summed the EEG data from matching audio-only and visual-only stimuli (i.e., audio-only and visual-only stimuli that came from the same original AV video; Fig. 1A). Thus, for each of the original 15 videos, we ended up with AV EEG responses and corresponding A+V EEG responses. Then, we used CCA to relate the multivariate speech representations (spectrogram + phonetic features) to each of these two EEG responses (AV and A+V). This provides two sets of rotation matrices, one between the stimulus and the AV EEG and one between the stimulus and the A+V EEG.

**Figure 1.**
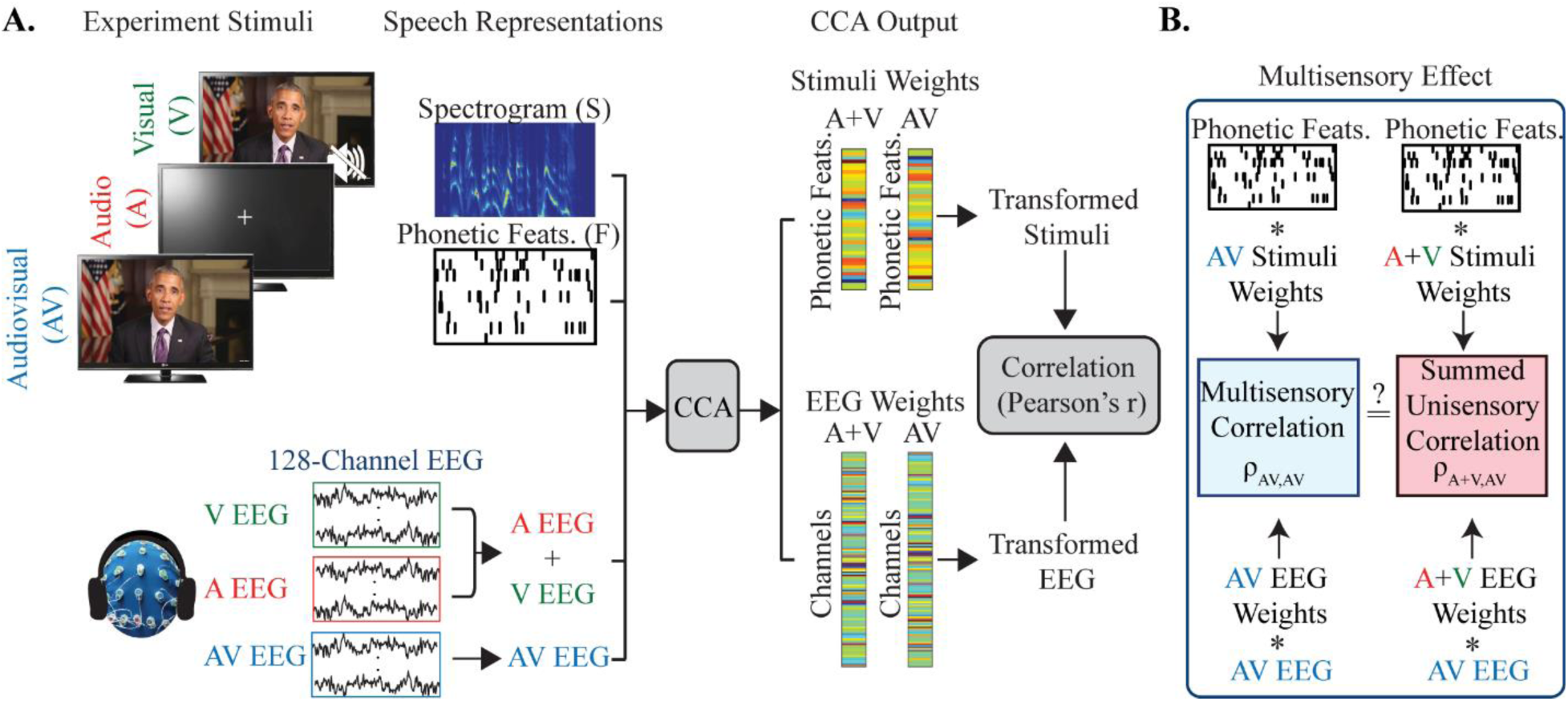
Experiment set-up and analysis approach. A. The stimulus representations used are the spectrogram and the phonetic features and are estimated directly from the speech stimuli. Below, the EEG recordings corresponding to each condition. The unisensory A and V EEG are summed to form an A+V EEG data set. The EEG and speech representations are used as inputs to the CCA in order to determine the optimum weights for rotating the EEG and the given stimulus representation for maximizing the correlation between the two. B. The model built using the A+V data is then tested on the left-out AV data in order to determine the presence of a multisensory effect.

Now, if the scalp recorded EEG activity for the AV condition is simply the auditory and visual modalities being processed separately with no integration occurring, then the A+V and AV EEG responses should be essentially identical. And we would then expect the rotation matrices learned on the A+V EEG data to be identical to those learned on the AV EEG data. Carrying this logic even further, we would then expect to see no differences in the canonical correlation values obtained from the AV data when using the CCA rotation matrices found by training on the A+V EEG data compared with the matrices found by training on the AV EEG data (Fig. 1B). In other words, we compared the correlation values obtained when we applied the AV weights (i.e., the A and B matrices found by training on AV EEG data) to left-out AV data, with the correlation values obtained when applying the A+V weights (i.e., the A and B matrices found by training on A+V EEG data) to the AV data (Fig. 1B). If there is some difference in the EEG response dynamics for multisensory (AV) compared with the summed unisensory activity (A+V) then we would expect this to have a significant effect on the canonical correlations since the A+V weights would not capture this whereas the AV weights would. To measure the size of this difference we calculate multisensory gain using the following equation:

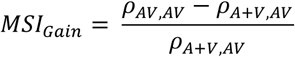

Where *ρ* is the canonical components correlation, the first subscript represents the rotations used and the second subscript represents the data on which those rotations are applied (Fig. 1B).

### Statistical analysis

All statistical comparisons were conducted using non-parametric permutation with 10,000 repetitions such that no assumptions were made about the sampling distribution (Combrisson and Jerbi, 2015). This was done by randomly assigning the values from the two groups being compared (pairwise for the paired tests, and non-pairwise for the unpaired tests) and calculating the difference between the groups. This process was repeated 10,000 in order to form a null distribution of the group difference. Then the tail of this empirical distribution is used to calculate the p-value for the actual data, and two-tailed tests are used throughout. Where multiple comparisons were carried out p-values were corrected using the False Discovery Rate (FDR) method (Benjamini and Hochberg, 1995). All numerical values are reported as mean ± SD.

## Results

### Robust indices of multisensory integration for the speech spectrogram and phonetic features

To investigate the encoding of more complex multivariate representations of the speech stimulus and to isolate measures of multisensory integration at different levels of the speech processing hierarchy we performed CCA on the AV EEG data using the spectrogram and phonetic features, having trained the CCA on (different) AV data and A+V data. We first sought to do this separately for the spectrogram representation and the phonetic representation to see if using either or both of these representations might show evidence of multisensory integration. And we also sought to do this for both our clean speech and noisy speech datasets.

In both conditions (clean and noisy speech) and for both representations (spectrogram and phonetic features) the correlations for the first component were significantly higher than for all other components. This suggests that the first component captures a substantial percentage of the influence of the speech on the EEG data. And, importantly, both representations showed evidence of multisensory integration.

For the spectrogram representation, we found significant multisensory effects (AV>A+V) for the first canonical component for speech in quiet (p<0.0001) and the first canonical component for speech in noise (p<0.0001, FDR corrected p-values; Fig. 2A, B). Indeed we found multisensory effects for 15/16 components for speech in quiet and for 15/16 components for speech in noise.

**Figure 2.**
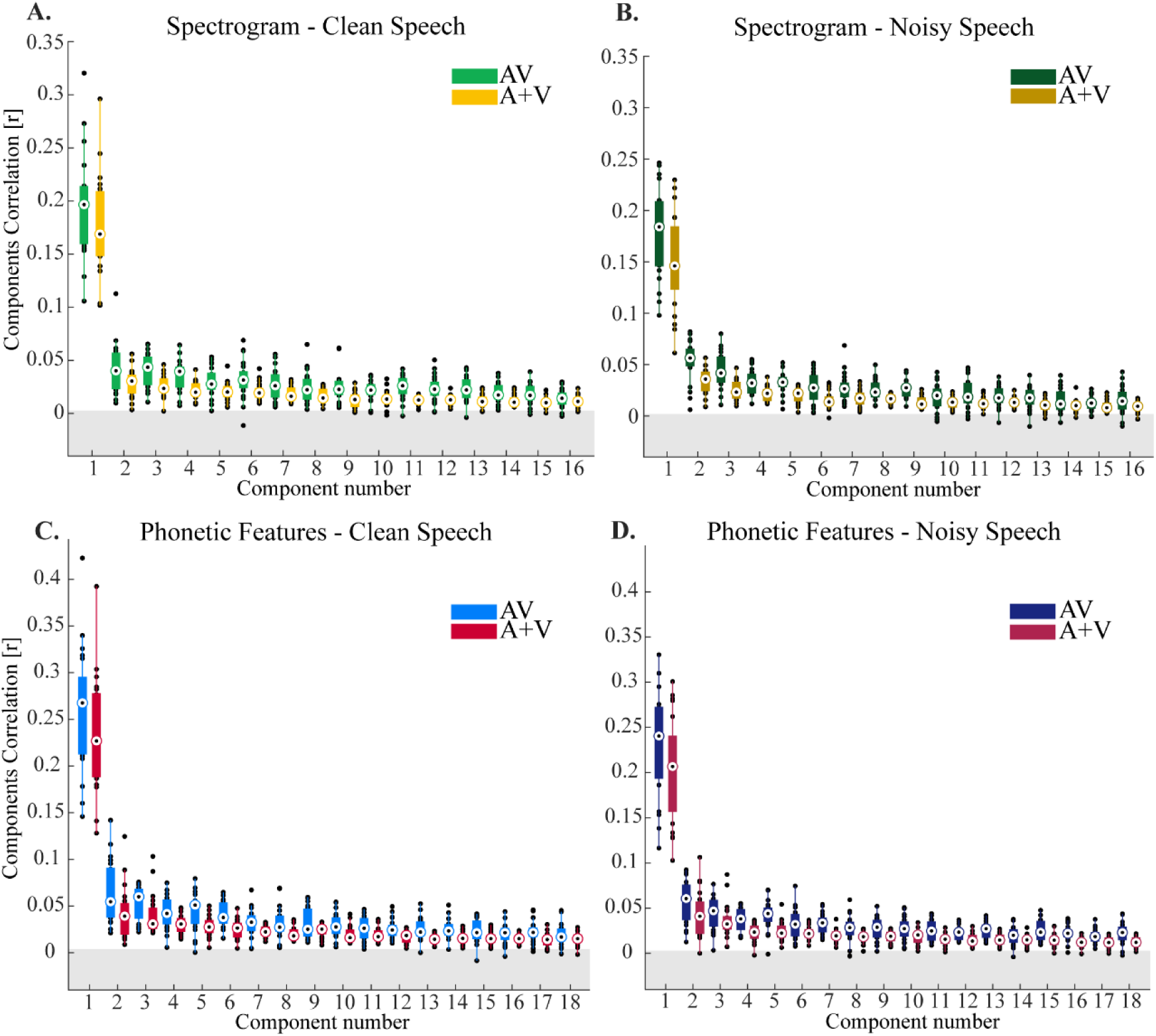
CCA analysis using the spectrogram and phonetic feature representation of the speech stimulus. A-B The canonical correlations for the spectrogram representation for speech in quiet and speech in noise respectively. C-D canonical correlations for the phonetic feature representation for speech in quiet and speech in noise respectively. The gray band represents an approximate chance level. All AV and A+V components performed above chance level p<0.0001 for speech in quiet, and for speech in noise p<0.0001.

A similar pattern was observed when examining the stimulus-EEG relationship using the phonetic feature representation of speech. Specifically, we also found multisensory effects for the first component for clean speech (p<0.0001) and for speech in noise (p<0.0001, FDR corrected). And we found multisensory effects for 13/18 components for speech in quiet and for all 18 components for speech in noise.

The fact that the spectrogram and phonetic feature representations produced qualitatively similar patterns of multisensory integration was not surprising. This is because both representations are mutually redundant; a particular phoneme will have a characteristic spectrotemporal signature. Indeed, if each utterance of a phoneme were spoken in precisely the same way every time, then the spectrogram and phonetic feature representations would be functionally identical (Di Liberto et al., 2015). But, as we discuss below, in natural speech different utterances of a particular phonemes will have different spectrograms. So, to identify the unique contribution of “higher-level” neurons that are invariant to these spectrotemporal differences and are sensitive to the categorical phonemic features we will need to index the EEG responses that are uniquely explained by the phonetic feature representation whilst controlling for the spectrogram representation. (Please see the section on **Isolating multisensory effects at the spectrotemporal and phonetic levels** below).

### Spatiotemporal analysis of canonical components: increased cross-modal temporal integration and possible increased role for visual cortex for speech in noise

The previous section showed clear evidence of multisensory integration in the component correlation values obtained from CCA. But how can we further investigate these CCA components to better understand the neurophysiological effects underlying these numbers? One way is to examine how these multisensory effects might vary as a function of the time-lag between the stimulus and EEG and how any effects at different time-lags might be represented across the scalp. This is very much analogous to examining the spatiotemporal characteristics of event-related potentials with EEG.

To investigate the spatiotemporal properties of the AV and A+V CCA models we ran the CCA at individual time-lags from −1s to 1.5s. We chose to focus our analysis on the first three canonical components. This was mostly to allow investigation of the dominant first component, but also to check whether or not useful insights might be gleaned from any of the subsequent components. For the first component of the spectrogram model, there was significant differences between AV and A+V at −300-(−200) ms, 0-125 ms and at 500-750 ms for speech in quiet (FDR corrected). For speech in noise there was significant differences at time shifts of −650-(−250) ms, −60-750 ms (Fig. 3A, D, FDR corrected). For the second and third components there was no clear pattern to the single time-lag correlation as it was quite flat across all time shifts for both clean and noisy speech (Fig. 3B, C, E, and F).

**Figure 3.**
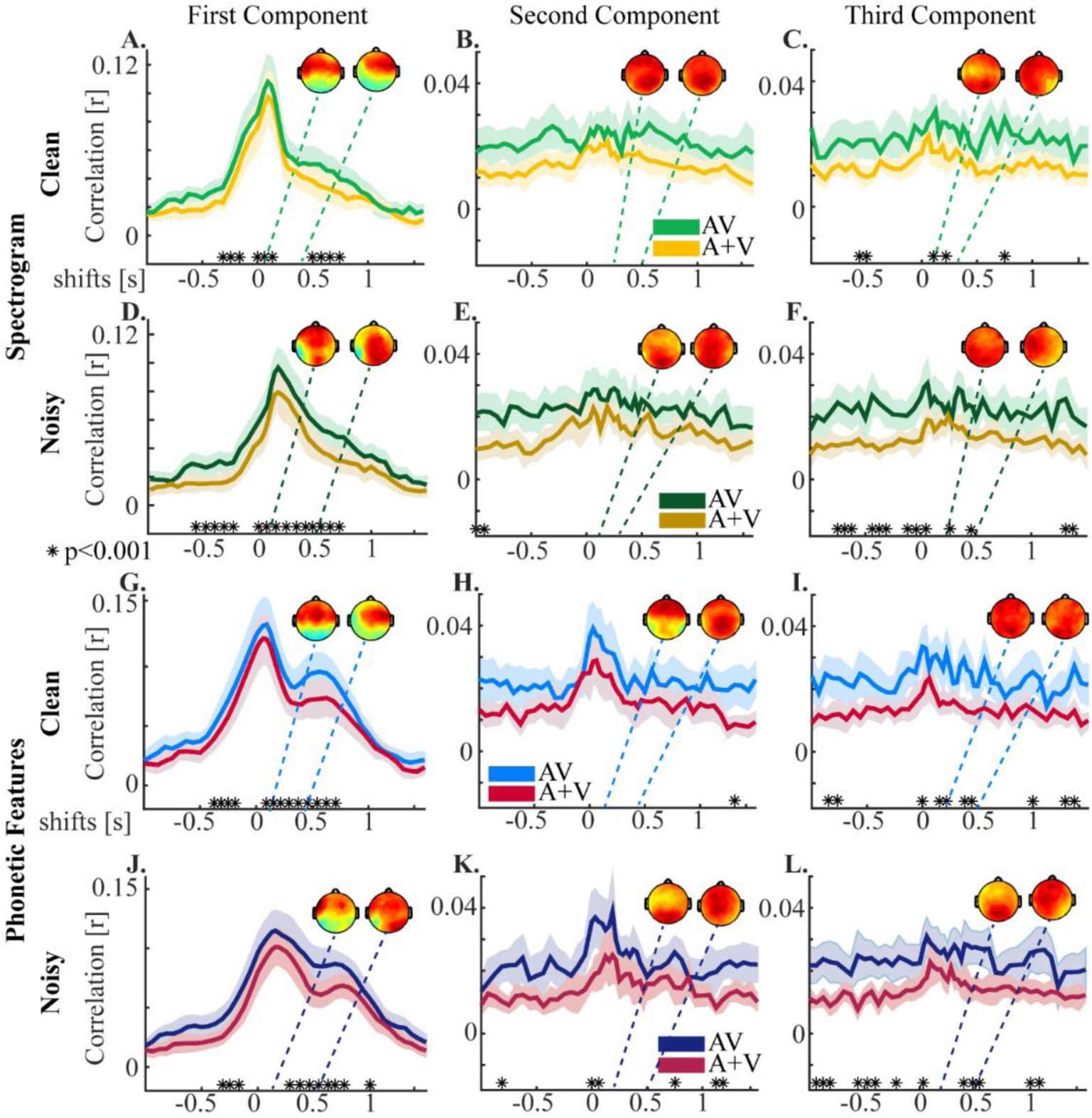
Correlations for the first three canonical components using the spectrogram and phonetic feature representation of the speech stimulus at single time shifts. A-C. The correlations using the spectrogram representation for the first three components using the AV model and the A+V model for speech in quiet, and D-F for speech in noise. G-I The single time shift correlations for the phonetic feature representation between the first three components for the AV and A+V model with the original raw AV EEG data for speech in quiet, and J-I for speech in noise. The respective topographies inset show the corresponding spatial correlations between the corresponding component of the AV model and the AV EEG. *p<0.001 FDR corrected.

For the phonetic feature representation we found significant differences between AV and A+V at −370-(−125) ms, 90-750 ms for speech in quiet and for speech in noise there was significant differences at time shifts of −300-(−125) ms and 300-750 ms (Fig. 3G, J, FDR corrected). For the second component the pattern reflected that of an onset response and while there was no difference between AV and A+V for speech in quiet there was a small window of difference for speech in noise at 0-75ms (Fig. 3H, K, FDR corrected). There was no clear pattern to the single time-lag correlation for the third component for both clean and noisy speech (Fig. 3I, L). In general, the number of lags at which there was a significant difference between AV and A+V was greater for noisy speech than clean speech, which is consistent with findings in Crosse et al. (2016b). The finding of significant differences at negative lags may be due to the fact that the EEG data is time-locked to the onset of the audio and since the visual information often precedes the audio (Schwartz and Savariaux, 2014; Chandrasekaran et al., 2009), the AV data may contain some information about the speech at ‘negative’ time-lags. Another possibility however, is that the effect at negative lags is due to the autocorrelation of the stimulus and the autocorrelation of the EEG. Nonetheless, in our multi-lag CCA we have only used positive lags (0-500 ms) and so any effects at negative lags will not influence our overall results.

To visualize the scalp regions underlying these components we calculated the correlation coefficient between each component from the AV models for each time lag with each scalp electrode of the AV EEG. This gives a sense of how strongly the data on each channel has contributed to that component. The spatial pattern for the first component revealed strong contributions from channels over central and temporal scalp for speech in quiet for both the spectrogram and phonetic feature representations. For speech in noise there was an additional correlation with occipital regions, possibly indicating an increased contribution from visual areas to multisensory speech processing in noisy conditions. Occipital channels also made clear contributions for the second and third components for both conditions, however due to the lack of a clear temporal response for these components, we are hesitant to over-interpret this.

### Isolating multisensory effects at the spectrotemporal and phonetic levels

As discussed above, the spectrogram and phonetic feature representations are highly correlated with each other. As such, measures of how well each individual representation maps to the EEG (as in Fig. 2) are difficult to interpret in terms of multisensory effects at specific hierarchical levels. To pinpoint effects at each specific level, we need to identify the unique contributions of the spectrogram and the phonetic feature representations to the EEG data. To do this, we first regressed out the spectrogram representation from the EEG and then related the residual EEG to the phonetic features using CCA (Bednar and Lalor, 2020). This should isolate the unique contribution (if any) provided by the phonetic feature representation. Examining such a measure across our 2 conditions (clean speech and noisy speech) allowed us to test the hypothesis that multisensory integration effects should be particularly pronounced at the phonetic feature level for speech in noise. We also performed a similar analysis for the spectrogram representation to test its unique contribution to the multisensory effect in quiet and noise. In this case, we regressed out the phonetic feature representation from the EEG and then related the residual EEG to the spectrogram using CCA. Again, we did this for both speech in quiet and speech in noise, to test for interaction effects on our multisensory integration measures between acoustic and articulatory representations and speech in quiet and noise.

We limited our analysis here to the first canonical component due to its dominant role in the above results, as well as to the fact that it displays a greater consistency across subjects relative to the other components (please see next section of results and Fig. 6).

We found that multisensory gain at the level of acoustic processing (unique contribution from spectrogram) was significantly greater than zero for both clean speech and noisy speech (Fig. 4A). However, there was no difference in this measure between conditions (Fig 4B; p = 0.92), suggesting that multisensory integration at the earliest cortical stages was similar for speech in quiet and noise. Meanwhile, multisensory gain at the level of articulatory processing (unique contribution from phonetic features) was also significantly greater than zero for both clean speech and speech in noise (Fig 4C). Importantly however, in line with our original hypothesis, there was a significant difference in this measure between conditions, with MSI gain being larger for speech in noise than speech in quiet (p = 0.02). This supports the idea that, when speech is noisy, the impact of complementary visual articulatory information on phonetic feature encoding is specifically enhanced.

**Figure 4.**
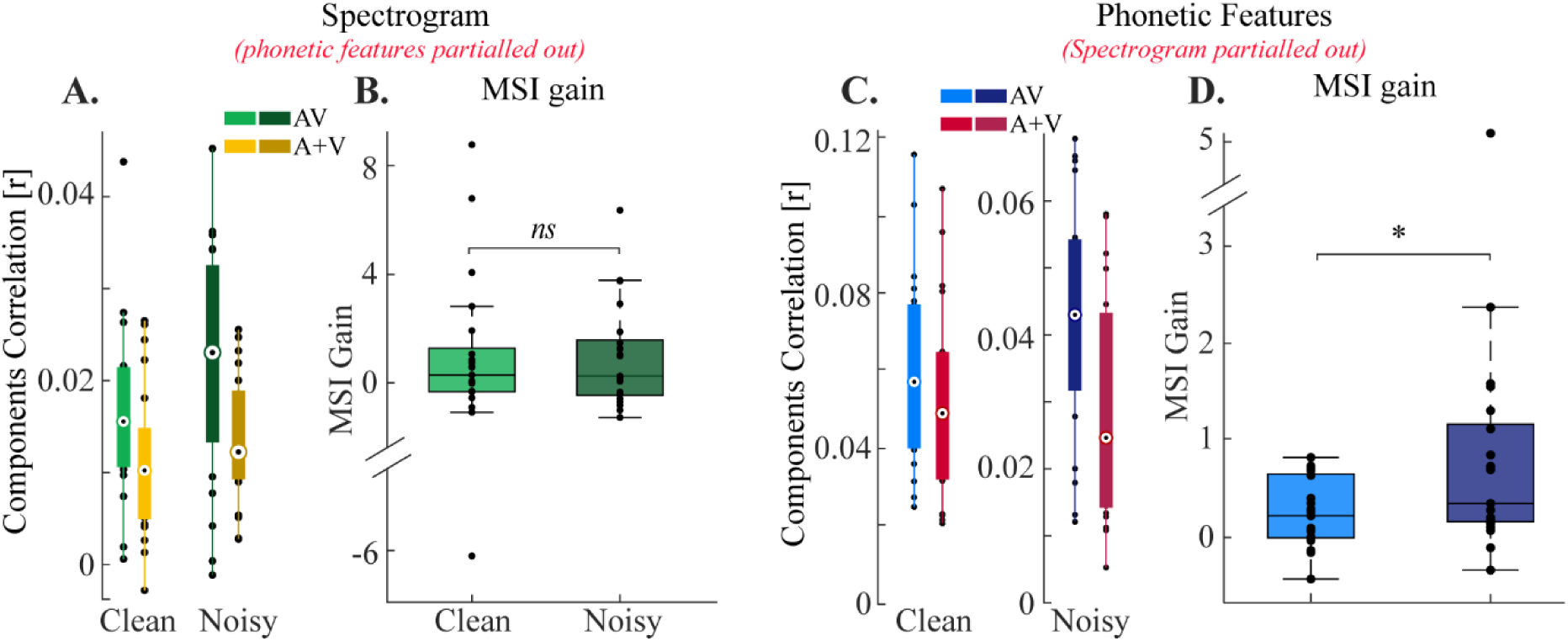
Multisensory gain for different speech representations for speech in quiet and in noise. A, B. For the unique spectrogram representation (phonetic features partialled out) there is no difference in gain, p=0.92. C, D. For the unique phonetic features (spectrogram partialled out) we find a difference in gain between conditions, p=0.02 (Unpaired permutation test for both).

We then wanted to examine whether the multisensory integration effects at the phonetic feature level might be driven by specific phonemes. More precisely, we wondered if the effect might be primarily driven by phonemes whose accompanying visual articulations are particularly informative (e.g., /b/, /p/ or /f/ compared to /g/ or /k/). To do this, we tested which phonemes were most correlated with the first canonical component. This involved taking the first component arising from the rotation of the phonetic feature stimulus on the left-out data and then calculating the correlation between this component and the time-series of each phoneme. If there is a high correlation between the component and a particular phoneme then it suggests that this phoneme is strongly represented in that component and it plays a role in driving the stimulus-EEG correlations that we have reported here. This analysis revealed that phonemes such as /p/, /f/, /w/, and /s/ were most strongly represented in the first component. In general, it also showed that consonants were more correlated with the component than vowels (Fig. 5A, B), although for speech in noise this effect was slightly less pronounced (Fig. 5B). To see these results in terms of visual articulatory features, we grouped the phonemes into visemes (the visual analog of phonemes), based on the mapping defined in (Auer Jr and Bernstein, 1997). This showed that bilabials (/b/,/p/ and /m/) and labio-dentals (/f/, and /v/) were most the features most correlated with the first component. Finally, we also checked that these phoneme-component correlations were not simply explainable as a function of the number of occurrences of each phoneme. To check this, we tested for a relationship between the phoneme-component correlations and the number of occurrences of each phoneme (Fig. 5B grey line). No correlation between the two was found for either speech in quiet (p = 0.99) or speech in noise (p = 0.71).

**Figure 5.**
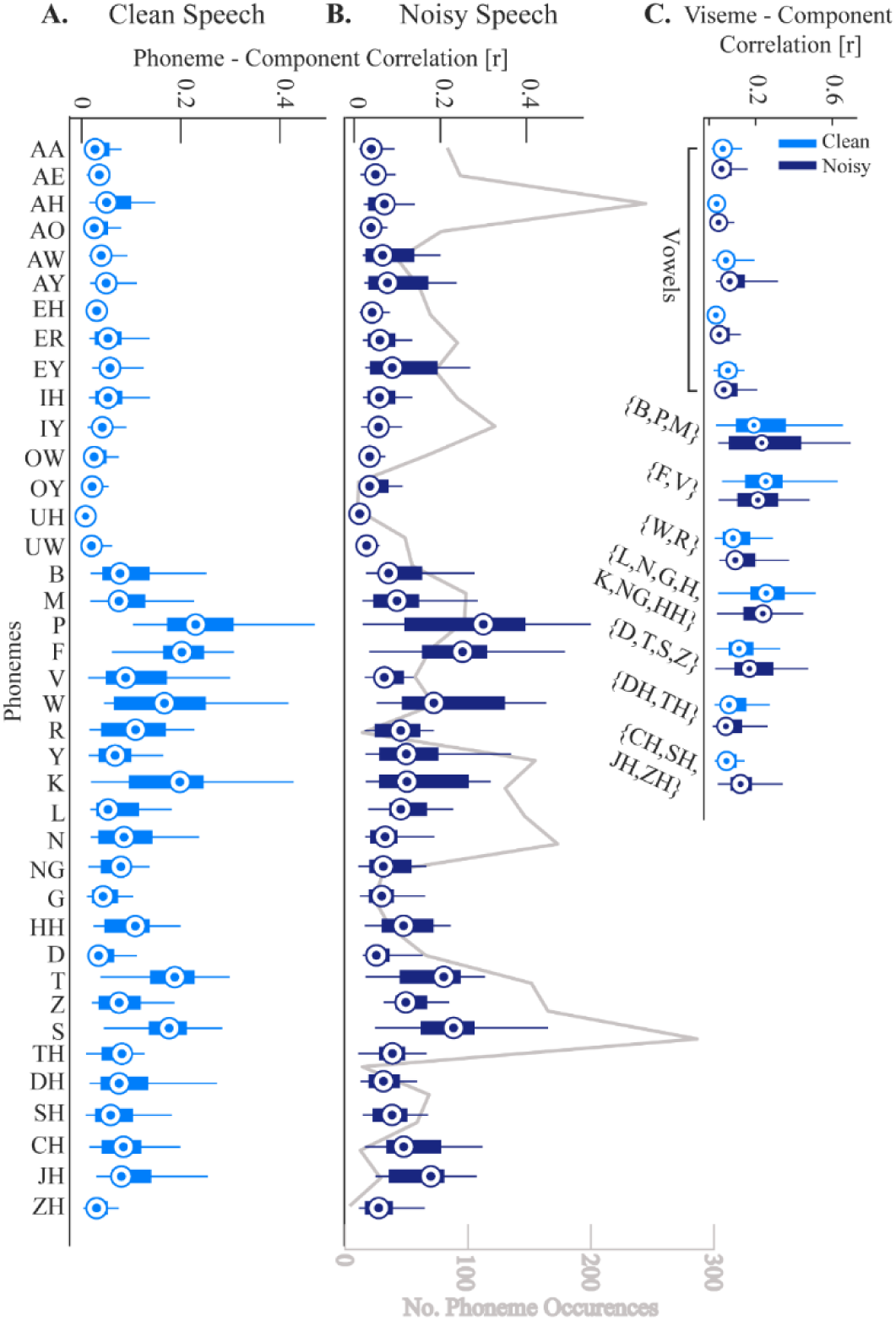
Correlation between phonemes and visemes time-series with the first canonical component. A. The correlations for each phoneme from the clean speech data. B. The correlations for each phoneme from the speech in noise data. The grey line plots the number of phoneme occurrences to show that it is not the case that the most frequent phonemes dominate the data. C. The correlation between each viseme (groups of visually similar phonemes) and the first component.

### Consistency of canonical components across subjects

CCA finds stimulus-EEG matrix rotations on a single subject basis. As such, for us to make general conclusions about results gleaned from individual canonical components, we must examine how similar the individual components are across subjects. To do this, we took the components for each subject and calculated the correlation (using Pearson’s r) for every subject pair (Fig. 6). Given its dominant role in capturing EEG responses to speech, and in the results we have presented above, we were particularly interested in consistency of component one across subjects.

**Figure 6.**
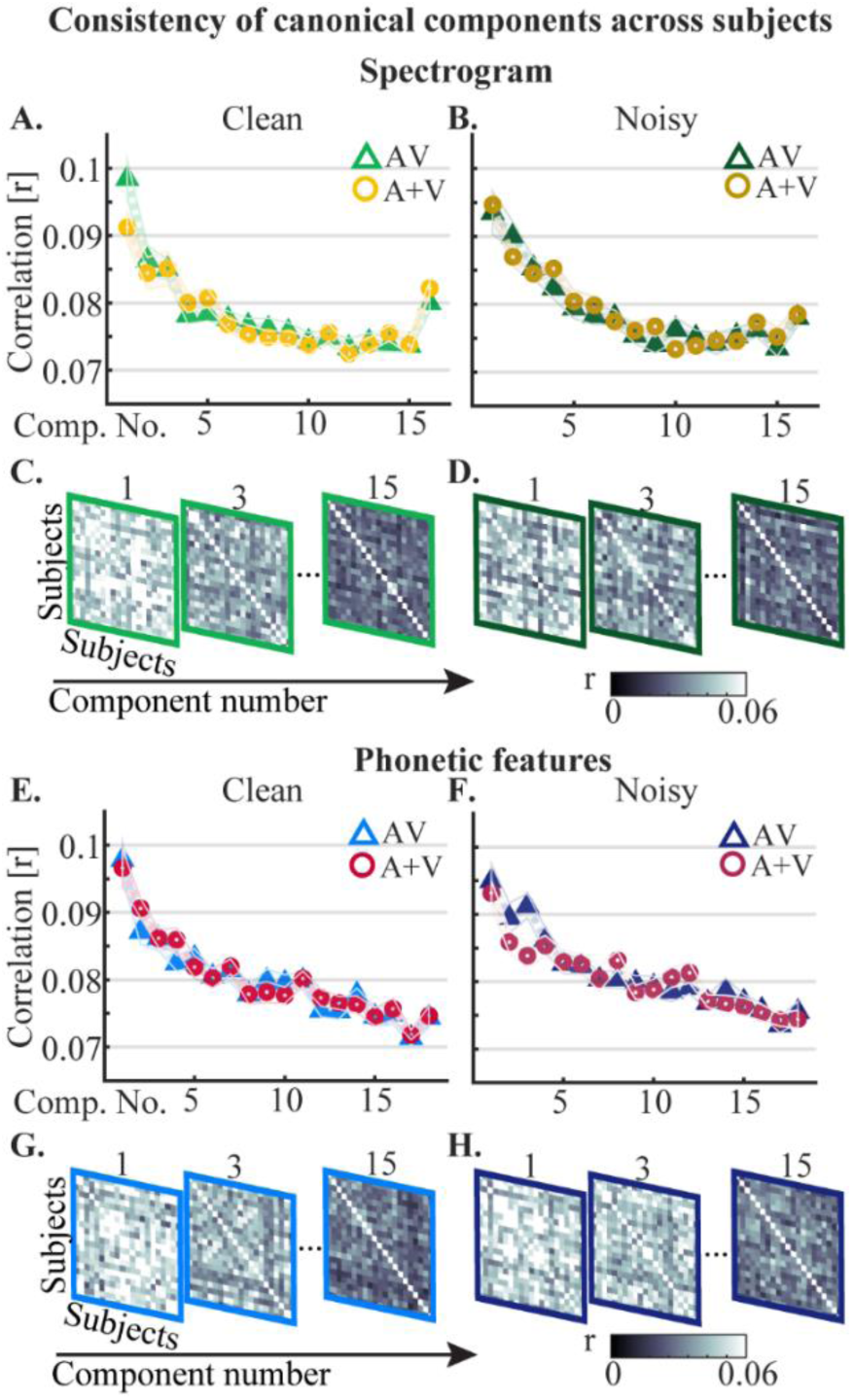
Consistency of canonical components across subjects for the spectrogram and the phonetic features after partialling out the other representation. A, B. Correlations of AV and A+V canonical components across subjects for the spectrogram representation for speech in quiet and speech in noise respectively. C, D. Correlation matrices visualizing the reduction in consistency in the AV component activity across subjects as the component number increases for the spectrogram. E, F. The correlation of the canonical components for the phonetic feature representation across subjects, and G, H. the corresponding correlation matrices for speech in quiet and speech in noise respectively.

For the spectrogram, the first components of the AV and A+V models were significantly more correlated across subjects than all other components for clean speech (p<0.0001 for both). For speech in noise the first component of the AV model was not significantly more correlated across subjects than the second component (p=0.055) but it had a significantly higher correlation than the remainder of the components (p<0.0001). The first A+V component for speech in noise was significantly more correlated across subjects than all others (p<0.001).

For the phonetic features model, a similar pattern emerged for the clean speech condition with the first components of the AV and A+V models being significantly more correlated across subjects than all other components (p<0.0001 for both). For noisy speech, the first component of the AV model was again significantly better than all others (p<0.02) and similarly the first component of the A+V model had a higher correlation than all others (p<0.0001).

Altogether, we found that the first component is most correlated across subjects, while later components are less correlated. This suggests that the first component, which dominated our results above, is also the most consistent component across subjects and, thus, that it is capturing processes that are general across those subjects. In contrast, the later components, as well as being smaller, are more variable across subjects, and, accordingly, may not be capturing similar underlying processes. This in turn can make results from these later components more difficult to interpret.

## Discussion

In this work, we have used a CCA-based framework for relating multivariate stimulus representations to multivariate neural data in order to study the neurophysiological encoding of multidimensional acoustic and linguistic features of speech. Our results show significant audiovisual integration effects on the encoding of the spectrogram and phonetic features of both clean and noisy speech. Importantly, these multisensory effects are enhanced at the phonetic level for speech in noise, supporting the hypothesis that listeners increasingly rely on visual articulatory information when speech is noisy.

### Enhanced multisensory integration effects at the phonetic-level of processing for speech in noise

It is well known that the enhancement of auditory speech processing provided by visual speech varies with listening conditions (Ross et al., 2007). However, the details of how visual speech impacts auditory speech processing at different hierarchical levels remains to be fully elucidated. There is a growing body of evidence indicating that AV speech integration likely occurs over multiple stages (Peelle and Sommers, 2015). In particular, it is thought that visual speech provides temporal information about the acoustic speech which can affect the sensitivity of auditory cortex (Grant and Seitz, 2000; Okada et al., 2013), as well as provide complementary cues that contain articulatory information which may be integrated with acoustic information in superior temporal sulcus (STS) (Beauchamp et al., 2004; Kayser and Logothetis, 2009; Nath and Beauchamp, 2011).

In this study, we found that audiovisual speech integration operates differently under different listening conditions (clean vs noisy (−9dB) speech). Specifically, for the encoding of low-level spectrogram features, we found that the integration effects are substantial for both speech in quiet and speech in noise. These integration effects are likely to be primarily driven by modulations of responses in early auditory cortex by temporal information provided by visual speech, which often precedes the auditory speech (Schwartz and Savariaux, 2014; Chandrasekaran et al., 2009). This result is also in line with recent work demonstrating multisensory benefits at the spectrotemporal level elicited by a visual stimulus that did not contain articulatory detail – dissociating the effect from access to higher-level articulatory details (Plass et al., 2019). Furthermore, the lack of any difference in the magnitude of these integration effects between clean and noisy speech conditions suggests that the benefits of visual speech provided at a low-level of processing might be similar regardless of acoustic conditions.

In contrast with this, using a higher-level phonetic feature representation, we found that the AV integration effects are different depending on the acoustic conditions (after regressing out the contribution of the spectrogram). Specifically, we found significantly larger integration effects for phonetic feature encoding in noisy speech than in clean speech. We suggest that this benefit is likely to be driven by an increased reliance on the visual articulations which help the listener to understand the noisy speech content by constraining phoneme identity (Karas et al., 2019). In line with this, we also show that the phonemes that most contribute to these results are those that have particularly informative visual articulations (Fig. 5).

While recent research has challenged the notion that scalp recorded responses to speech reflect processing at the level of phonemes (Daube et al., 2019), our findings reveal a sharp dissociation in AV integration effects on isolated measures of acoustic and phonetic processing across listening conditions. This seems difficult to explain based on considering acoustic features alone and seems consistent with the idea of visual articulations influencing the categorization of phonemes (Holt and Lotto, 2010). More generally, we take this as a further contribution to a growing body of evidence for phonological representations in cortical recordings to naturalistic speech (Brodbeck et al., 2018; Di Liberto et al., 2015; Gwilliams et al., 2020; Khalighinejad et al., 2017; Yi et al., 2019).

One brain region likely involved in exploiting the articulatory information when the speech signal is noisy is superior temporal sulcus (STS) which has been shown to have increased connectivity with visual cortex in noisy compared with quiet acoustic conditions (Nath and Beauchamp, 2011). While it remains an open question as to how much speech-specific processing is performed by visual cortex (Bernstein and Liebenthal, 2014), there is a some early evidence supporting the notion that visual cortex might processes speech at the level of categorical linguistic (i.e., phonological) units (O’Sullivan et al., 2017; Hauswald et al., 2018). If true, visual cortex would be in a position to relay such categorical, linguistic information to directly constrain phoneme identity, again, possibly in STS. On top of this it has been shown that frontal cortex selectively enhances processing of the lips during silent speech compared with when the auditory speech is present, suggesting an important role for visual cortex in extracting articulatory information from visual speech cues (Ozker et al., 2018). Thus it is plausible that the greater multisensory gain seen here for phonetic features when the speech is noisy is underpinned by an enhancement of mouth processing in visual cortex which feeds information about the articulations to STS where they influence the online processing of the acoustic speech.

### Investigating hierarchical stages of speech processing – CCA captures relationships between multi-dimensional stimuli and EEG

The event related potential (ERP) technique has for a long time been used to advance our understanding of the multisensory integration of speech (van Wassenhove et al., 2005; Shahin et al., 2018; Klucharev et al., 2003; Bernstein et al., 2008; Molholm et al., 2002; Meredith and Stein, 1993; Saint-Amour et al., 2007). However, this approach is ill suited for use with natural, continuous speech stimuli.

More recently, researchers have begun to use methods such as multivariate regression (Crosse et al., 2016a) in the forward direction (predicting neural data from the stimulus, Di Liberto et al., 2015; O’Sullivan et al., 2017; Lalor and Foxe, 2010; Ding and Simon, 2012; Golumbic et al., 2013; Broderick et al., 2018) and backward direction (stimulus reconstruction from neural data, Crosse et al., 2015a; Crosse et al., 2016b; Crosse et al., 2015b; Mesgarani et al., 2009), which allows characterization of neural responses to natural speech. However, regression models in their general form allow only univariate-multivariate comparison, whereas with CCA one can relate multivariate stimulus representations (discrete/continuous) to multivariate neural responses. This is a useful advance over current techniques to study speech processing since CCA can use all features (of the stimulus and the neural response data) simultaneously to maximize the correlation between the speech representation and the neural data (de Cheveigne et al., 2018). Importantly, this approach has allowed us to answer questions which we could not do with previous methods, such as the impact of visual speech on auditory speech processing at different stages.

Our results show significant multisensory interaction effects in EEG responses based on the spectrogram and phonetic feature representations of the speech signal and so provides support for the multistage framework for audiovisual speech integration. Examining the relationship between the stimulus representations and EEG data at individual time-shifts reveals a peak in the correlation at around 100 ms post-stimulus for both the spectrogram and phonetic feature representations. This is likely attributable to a sound onset response. For the phonetic feature representation however, there is also a second broad peak at around 300-600 ms whereas for the spectrogram there is no noticeable second peak. In terms of the scalp regions which most contribute to the first component, we found it to be dominated by central and temporal regions for speech in quiet, and for speech in noise there is a greater contribution from more parietal and occipital regions. This is likely due to increased contributions from the visual areas when the acoustic speech is noisy.

### Conclusion

This work has used a novel framework to study multisensory interactions at the acoustic and phonetic levels of speech processing. This has revealed that multisensory effects are present for both the spectrogram and phonetic feature representations when the speech is in quiet or when it is masked by noise. However, for speech in noise, multisensory interactions are significantly larger at the phonetic-feature level suggesting that in noisy conditions, the listener relies more on higher-level articulatory information from visual speech.

